# Retrospective attention reveals a decaying theta rhythm in conscious access to a preceding stimulus

**DOI:** 10.1101/2025.09.23.677981

**Authors:** Sugeun Yun, Younglae Kim, Jaeseung Jeong, Yee Joon Kim

## Abstract

Conscious perception is often presumed to arise immediately upon external stimulation. However, the phenomenon of retroperception-conscious access to a stimulus after it has disappeared-challenges this view by demonstrating that awareness can emerge retrospectively. The fine-grained temporal dynamics governing this process, however, remain unknown. Here, we tested three competing models for the temporal architecture of retroperception: a monotonic decay, a sustained oscillation, and a damped oscillation. We implemented an established retro-cue paradigm and densely sampled behavioral reports. Through spectral analysis and computational modeling, we revealed that both behavioral accuracy and subjective visibility exhibited damped oscillations in the theta-band, superimposed on a time-dependent decay. Furthermore, we demonstrated that these oscillations in accuracy and visibility were temporally aligned, advocating a functional interplay between attention and perception in the post-stimulus period. These results reveal a rhythmically structured and temporally constrained window for conscious reactivation, redefining the limits of when and how the mind can become aware of the past.

## Main

How and when sensory events enter awareness remain central questions in consciousness research^1–3^. Classic models of perception posit that awareness is tightly coupled to a stimulus presentation, typically emerging -100-200 ms after its onset^4–6^. *Retroperception*, conscious access to latent sensory traces after the stimulus has disappeared, challenges this view^7^. By decoupling the formation of awareness from stimulus presentation, retroperception reveals a temporal flexibility of conscious access and constrains the neural mechanisms that support it^8^.

In retroperception, a *retro-cue*, cue presented after target offset, can direct attention to the target’s location and trigger conscious access of the previously unnoticed target^7–11^. Recent electroencephalogram (EEG) research further underscores attention’s critical role in retroperception^12^. Yet, the fine-grained temporal dynamics of retroperception remain unknown, as previous studies have sampled at only a few cue-target intervals (CTIs). This gap persists even though an early dense-sampling study did examine the temporal dynamics of attentional sampling in both pre- and retro-cue conditions, speculating rhythmicity in both^13^. The rhythmic sampling was observed in the pre-cue conditions, with numerous studies confirming the dynamics in both attentional and perceptual sampling^14–19^. In contrast, the study failed to detect rhythmicity in retro-cue conditions, likely due to backward masking from the salient-cue. This has left the temporal dynamics of retroperception largely unknown. Clarifying the dynamics would define the temporal boundary of conscious access, unifying attentional sampling with retroperception.

To address this gap, we employed a briefly dimmed circle as a retro-cue to minimize masking effect and densely sampled behavioral reports across CTIs to resolve the temporal structure. Building on this design, we hypothesize three candidate temporal architectures of retroperception.

1. Monotonic Decay: The magnitude of retro-cue-induced enhancement of visual experience gradually decreases over time without any oscillation, producing a monotonic decay in subjective visibility across CTI as illustrated in Fig. 1a.
2. Sustained Oscillation with a Decaying Trend: Retroperception exhibits rhythmic oscillations superimposed on a decaying trend, resulting in fluctuation of subjective visibility across CTI as illustrated in Fig. 1b.
3. Damped Oscillation with a Decaying Trend: Retroperception oscillates rhythmically, but the amplitude progressively diminishes, eventually converging to a non-oscillatory state while following an overall decay. This would appear behaviorally as damped oscillations superimposed on a decaying trend of subjective visibility over CTI as illustrated in Fig. 1c.

**Fig. 1.**
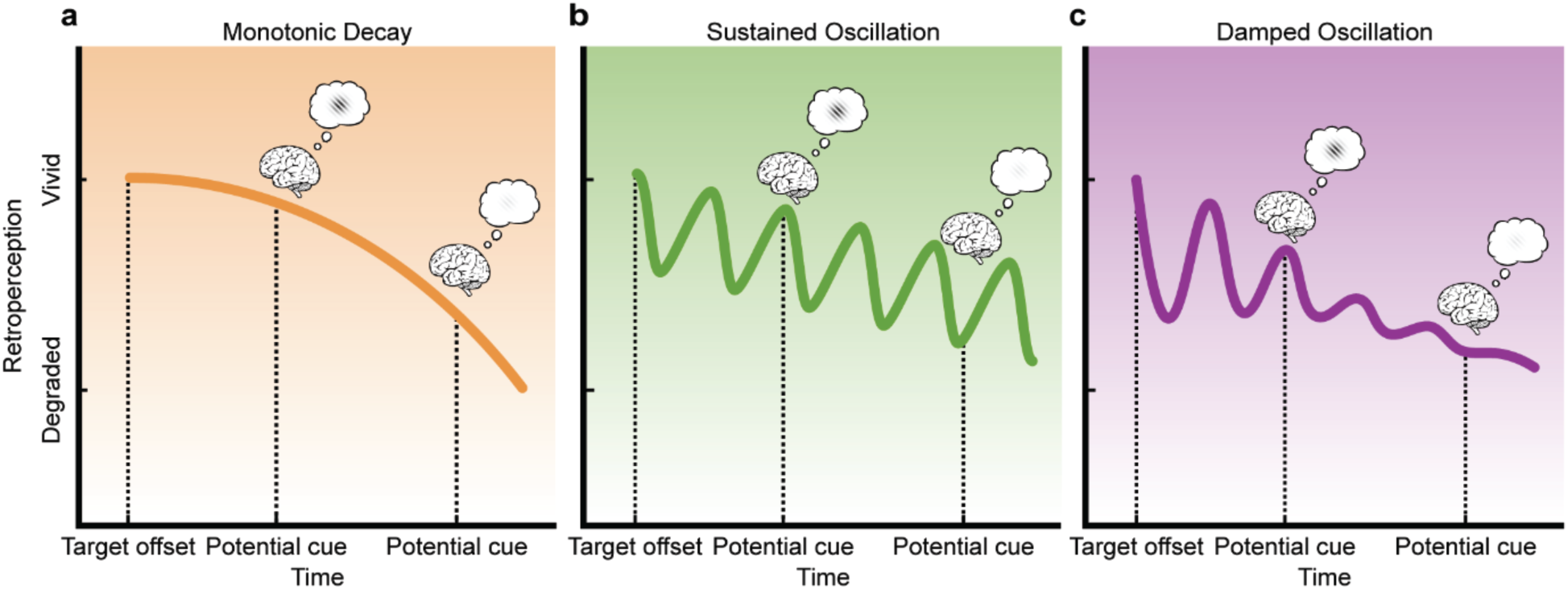
Schematic representation of hypotheses on the temporal dynamics of retroperception. The black dotted lines indicate the timing of key events (i.e., target offsets and cue presentations). Colored lines in each panel represent the hypothesized temporal dynamics of retroperception. When a cue appears at a favorable time point, participants perceive the target clearly, illustrated by bright Gabor patches within the brains. Conversely, when a cue appears at an unfavorable time point, participants experience degraded perception, illustrated by faded Gabor patches. **a** Monotonic decay model (orange line): Monotonic degradation of visual experience. **b** Sustained oscillation model (green line): Rhythmic oscillation superimposed on a decaying trend. **c** Damped oscillation model (purple line): Damped oscillation superimposed on a decaying trend.

To test these competing hypotheses, we employed a dense-sampling paradigm in which both discrimination accuracy and visibility rating measures were collected across a wide range of CTIs, spanning from the pre-cue to retro-cue condition. We then examined whether retroperception exhibits rhythmic fluctuations, and if so, whether these oscillations follow a sustained or damped profile. Temporal dynamics were analyzed using spectral decomposition with Irregular Resampling Auto-Spectral Analysis (IRASA) to isolate oscillatory components from fractal noise, followed by individualized model fitting to assess the relative explanatory power of each hypothesized pattern. Finally, we evaluated the phase alignment between accuracy and visibility measures to probe functional coupling between attention and perception. This comprehensive approach allowed us to characterize not only the presence of rhythmicity in retroperception, but also its temporal constraints and underlying structure.

## Result

The main experiment aimed to capture the temporal dynamics of target discrimination accuracy and subjective visibility of the target (Fig. 2a). Our experimental paradigm was adapted from the previous study^7^, except that we tested 146 pre-cue SOAs before target and 146 post-cue SOAs after target, for each of the left and right cueing conditions. At the beginning of each trial, participants were presented with a central fixation point flanked by two circular placeholders, aligned horizontally to the left and right of the fixation. A low-contrast Gabor patch, with a randomly determined orientation, served as the target stimulus and was presented for 50 ms within one of the two placeholders. The contrast level of the Gabor stimuli was individually adjusted for each participant prior to the main experiment to ensure consistent perceptual difficulty. A cue was presented by briefly dimming one of the placeholders to avoid masking the target, with 50% validity. The CTIs ranged from -1,500 ms to -50 ms in the pre-cue condition (Fig. 2b) and from 50 ms to 1,500 ms in the retro-cue condition (Fig. 2c), in 10 ms steps, and pre- and retro-cue trials occurred with equal probability to minimize expectation effects. Participants performed an orientation discrimination task and reported subjective visibility of the target at the end of each trial. In the orientation discrimination task, a grating matching the target’s orientation and the other grating with an orthogonal orientation were presented. Participants indicated which of the two gratings corresponded to the target orientation. On the visibility response screen, participants were also instructed to rate the subjective visibility of the target on a scale from 1 (Not seen) to 8 (Maximal visibility) (See details in Method section).

**Fig. 2.**
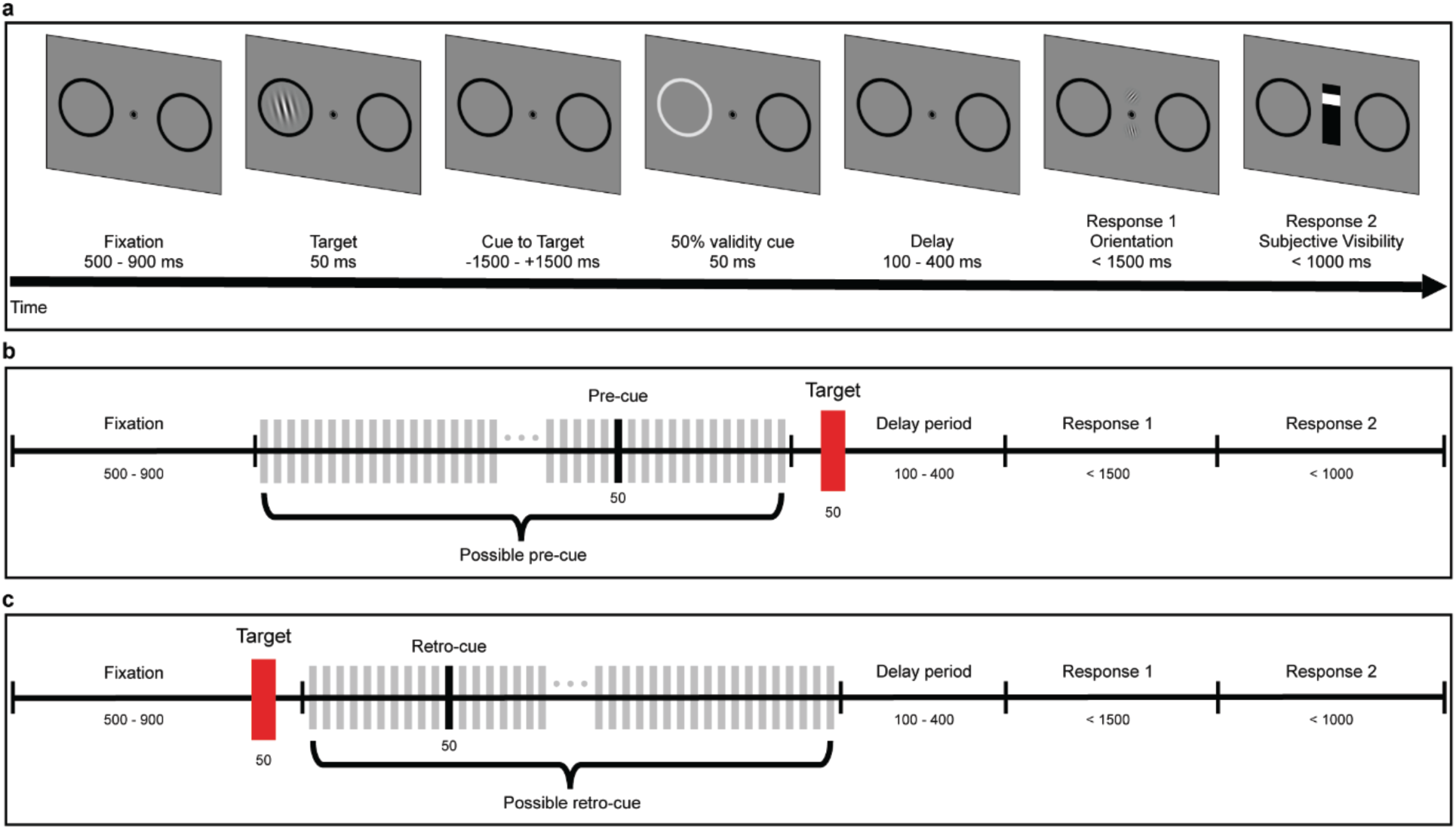
Experimental paradigm. **a** Example of trial structure. Each trial begins with a variable inter-trial interval (0.5-0.9 s), followed by the target presentation, appearing within one of the placeholders for 50 ms. A dimming of a placeholder, serving as a cue with 50% validity, is displayed for 50 ms. The cue-target interval (CTI) varies randomly from -1.5 s to 1.5 s. After a brief delay, participants report the identity and visibility of the target. **b** Example timeline of a pre-cue trial. The cue (black vertical line) is presented before the target onset (red vertical line). Possible cue onset times (gray vertical lines) range from -1.5 s to -50 ms relative to target onset, in 10 ms steps. Following the target offset, the response screen is presented after a brief delay. **c** Example timeline of a retro-cue trial. The cue (black vertical line) is presented after the target onset (red vertical line). Possible cue onset times (gray vertical lines) range from 50 ms to 1.5 s after target offset, in 10 ms steps. Following the cue offset, the response screen is displayed after a brief delay.

### Temporal dynamics of discrimination accuracy and subjective visibility

For each specific time point, behavioral accuracy was calculated by dividing the number of correct responses within a 100 ms time bin, centered on the time point, by the total number of trials within the bin. The average accuracy across all trials was approximately 80% (77.17 ± 1.36%; Symbol ± indicates the standard error). Similarly, visibility ratings were averaged within each 100 ms time bin to estimate subjective visibility at the corresponding time point, with an overall average visibility rating being approximately 4 (4.00 ± 0.20). To capture the temporal dynamics of accuracy and visibility, the time bins were shifted forward by 10 ms, and accuracy and visibility were calculated in the same manner. This process was repeated throughout all cue-to-target interval time points.

In retro-cue conditions, we observed alternating phases of increasing and decreasing behavioral accuracy (Fig. 3a & 3b). In addition to this wax-and-wane pattern, the behavioral accuracy as a function of a CTI showed signs of an overall decaying pattern in especially valid cue trials where target appeared at the cued placeholder (Blue graphs in the top rows of Fig. 3) and invalid cue trials where target appeared at the uncued placeholder (Red graphs in the top rows of Fig.3). Visibility ratings also exhibited a rhythmic fluctuation as well as gradual decay (Fig. 3c & 3d). Accuracy and visibility ratings in pre-cue conditions showed an overall increasing trend and rhythmic fluctuation, replicating findings from previous studies^13,14,16^ (Fig. S1).

**Fig. 3.**
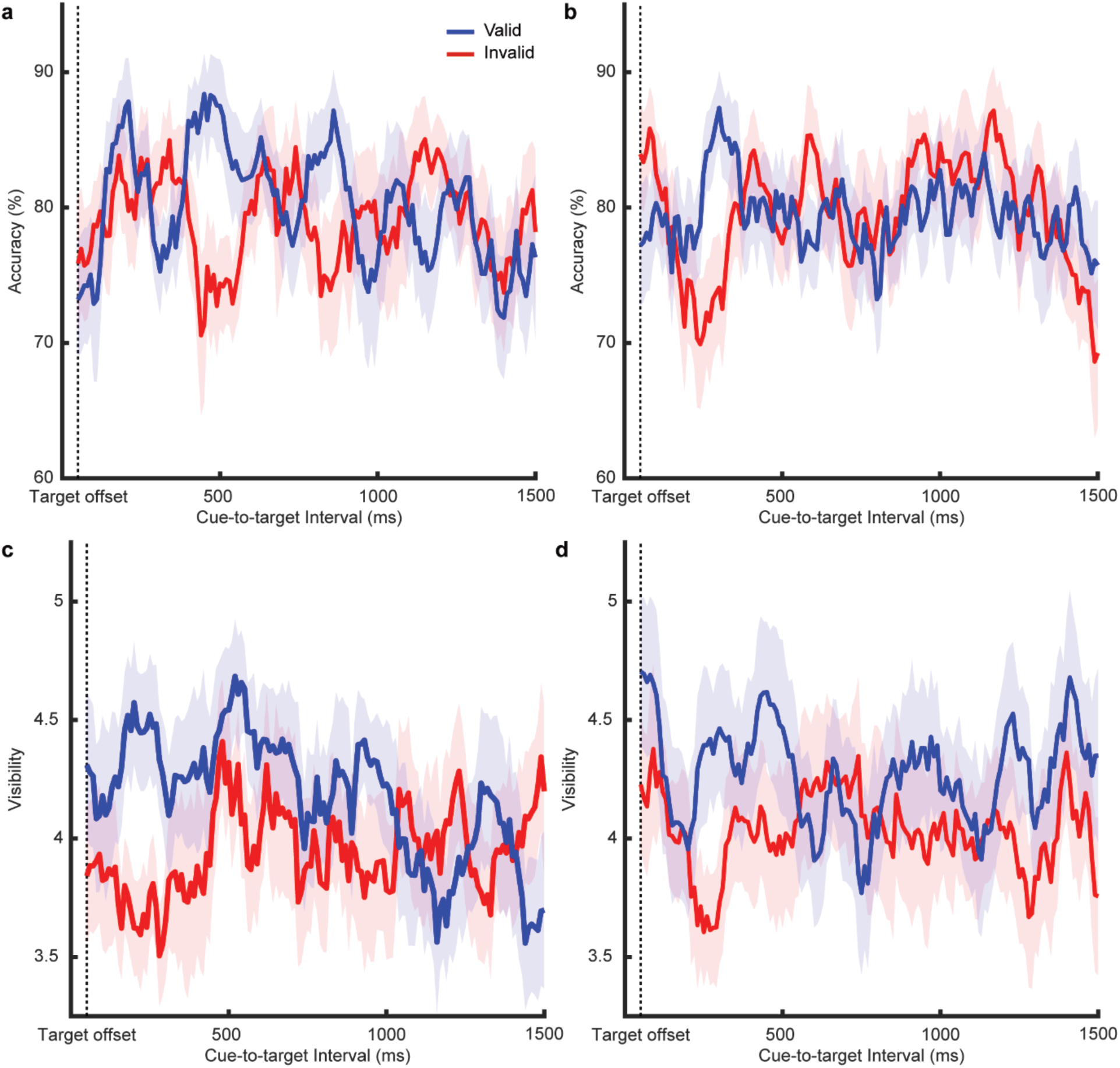
Temporal dynamics of discrimination accuracy and subjective visibility. Black dotted lines mark the time of the target offset. Blue lines and shaded areas correspond to valid cue trials; red lines and shaded areas correspond to invalid cue trials. Solid lines indicate the average performance (Top row) and visibility rating (Bottom row) across participants (n = 19). Shaded areas represent the standard error of the mean (SEM). **a** Target discrimination accuracy as a function of cue-target interval (CTI) for left-cued trials where the cue appeared in the left placeholder. **b** Target discrimination accuracy as a function of CTI for right-cued trials where the cue appeared in the right placeholder. **c** Subject visibility of the target as a function of CTI for left-cued trials. **d** Subjective visibility of the target as a function of CTI for right-cued trials.

Notably, although the overall trends in accuracy and visibility ratings differed between the pre-cue and retro-cue conditions, the observed rhythmic fluctuations in both cueing conditions indicate that oscillatory dynamics in visual information processing persist regardless of whether attention is cued before or after target presentation. Since the primary objective of the research is retroperception, we focus on the temporal dynamics of accuracy and visibility ratings in the retro-cue condition for further analysis.

### Rhythmicity in temporal dynamics of discrimination accuracy and subjective visibility

To determine whether both the behavioral accuracy and the visibility rating as a function of a CTI were rhythmic or mere random noise, we conducted a detailed spectral analysis using the established approaches in previous studies^16,20^. First, we applied IRASA to the observed time courses of accuracy and visibility ratings to isolate and analyze the rhythmic component from the 1/f background activity^20^. From the Fourier-transformed power spectrum of the original behavioral time course, we identified a prominent peak that exceeds the 1/f background activity as an oscillatory peak. We defined the center frequencies of the oscillatory peaks as individual peak frequencies (IPFs) for each cue condition and each participant separately. To identify the shared structure of the power spectrum across participants, we normalized and aligned the individual power spectra to each IPF (Fig. 4). The resulting average spectrum exhibited a single peak, indicating that the IPFs predominantly drive behavioral fluctuations. Regardless of valid or invalid trials in both left and right cueing conditions, we found that the behavioral time courses exhibited periodic fluctuations in the theta-band for both target discrimination accuracy (Fig. 4a & 4b) and visibility rating (Fig. 4c & 4d). These findings demonstrate that the fluctuations in the target discrimination accuracy are not random but reflect rhythmic oscillations at approximately 4 Hz, persisting even beyond target offset. At the same time, visibility ratings fluctuate rhythmically at similar frequencies, further suggesting the tight coupling between attention and perception.

**Fig. 4:**
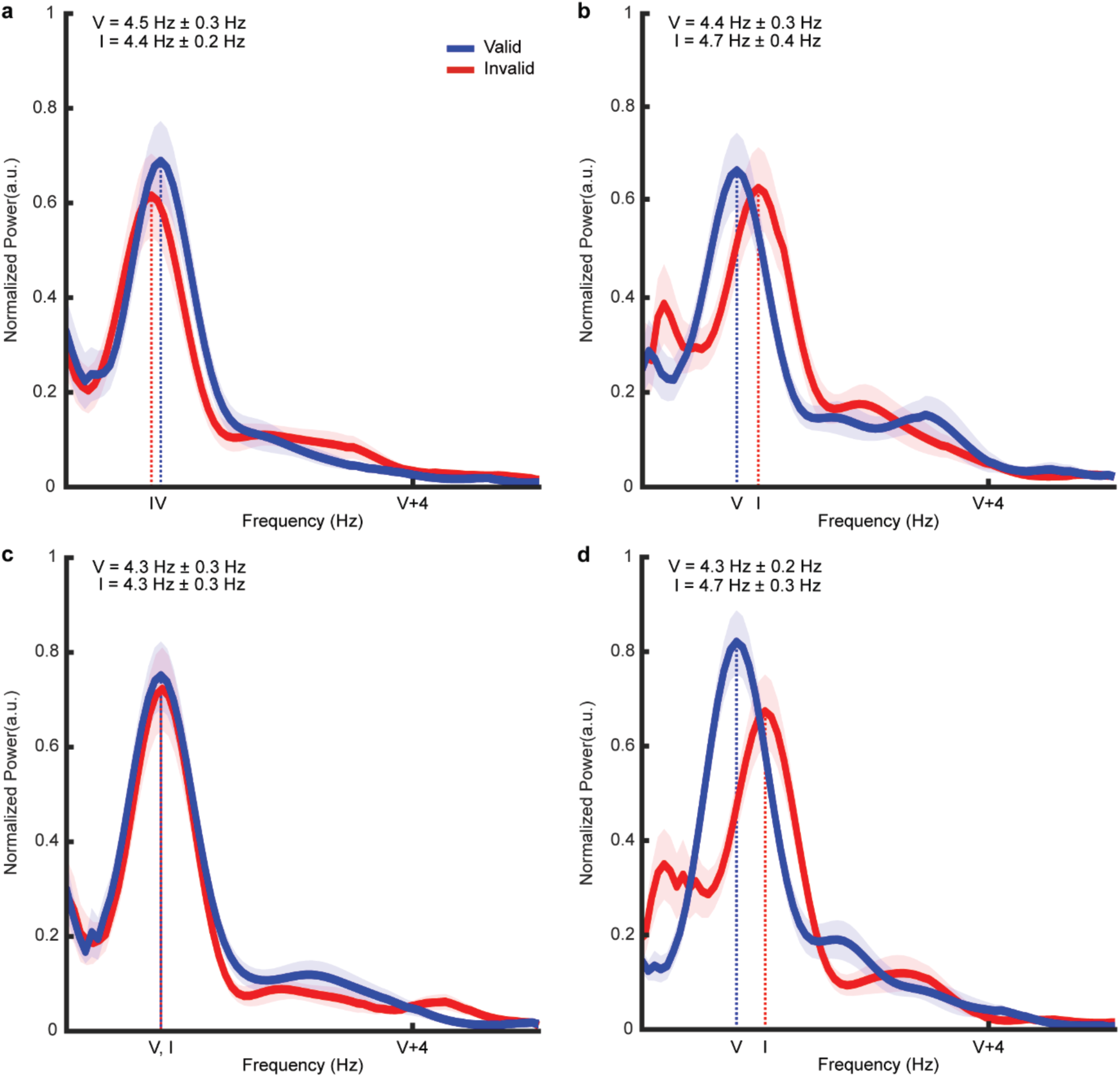
Power spectra of rhythmic oscillations. Blue (red) lines and shaded areas represent valid (invalid) trials. Solid lines indicate normalized power averaged across participants, shaded areas represent standard error of the mean (SEM), and dotted vertical lines mark the mean individual peak frequency (IPF). **a** Power spectra for target discrimination accuracy for left-cued trials. The mean IPF in valid trials was 4.5 ± 0.3 Hz, compared with 4.4 ± 0.2 Hz in invalid trials. **b** Power spectra for target discrimination accuracy for right-cued trials. The mean IPF in valid trials was 4.4 ± 0.3 Hz, compared with 4.7 ± 0.4 Hz in invalid trials. **c** Power spectra for the visibility ratings for left-cued trials. The mean IPF in valid trials was 4.3 ± 0.3 H, compared with 4.3 ± 0.3 Hz in invalid trials. **d** Power spectra for the visibility ratings for right-cued trials. The mean IPF in valid trials was 4.3 ± 0.2 Hz, compared with 4.7 ± 0.3 Hz in invalid trials.

### Model comparison: Testing hypotheses on retroperception

Given that the identified IPFs represent the most dominant frequencies in each participant’s behavioral time courses, we performed model fitting incorporating these IPFs to characterize the temporal profile of the retroperception. This analysis employed three distinct models derived from our theoretical assumptions. The first model is described as a quadratic function without any oscillatory component, where t represents the CTI, a_1_ and b_1_ are the coefficient of the quadratic and linear term, respectively (Equation (1)). This monotonic decay model corresponds to our first hypothesis, which states that retroperception decays over time without rhythmic fluctuation (Fig. 5a).

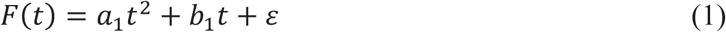

**Fig. 5:**
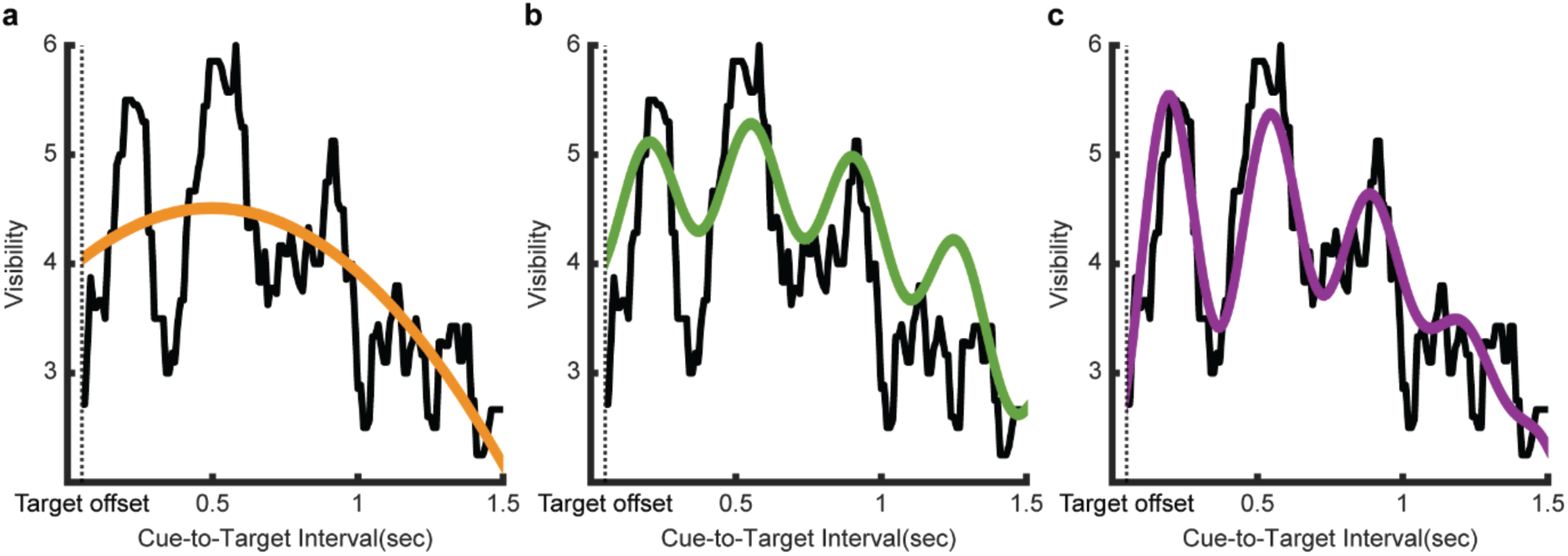
Model fits to a representative participant’s visibility time course. Black dotted lines mark the time of the target offset. Solid black lines represent a representative participant’s observed visibility rating fluctuations as a function of the cue-target interval. The three hypothesized models were fitted to this time course. a The orange line illustrates the fit of the monotonic decay model (Takeuchi’s information criterion [TIC] = 46.1) b The green line illustrates the fit of the sustained oscillation model (TIC = 35.2). c The purple line illustrates the fit of the damped oscillation model (TIC = 28.8).

The second model adds a first-order Fourier series to the first model, using the IPF determined by IRASA analysis. In the second model, f is the IPF for the Fourier series a_2_ and b_2_ are the amplitudes for cosine and sine components, respectively. (Equation (2)). This sustained oscillation model reflects our second hypothesis, positing that retroperception exhibits similar rhythmic oscillation as observed in the pre-cue conditions, along with a monotonically decaying trend (Fig. 5b)

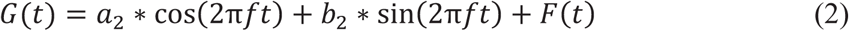

The third model introduces a linear damping parameter to the amplitude of the first-order Fourier series. In the third model, f is the IPF for the Fourier series, a_3_ and b_3_ are the amplitudes for the cosine and sine components, respectively, and il_1_and il_2_ are the linear damping coefficients applied to the amplitudes (Equation (3)). This damped oscillation model corresponds to our third hypothesis, which proposes that retroperception exhibits a damping oscillation with an overall decaying trend (Fig. 5c).

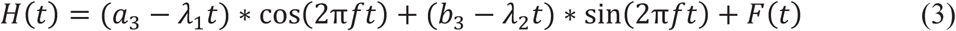

We fitted each model to the individual time course of accuracy and visibility rating using the maximum likelihood estimation. To account for potential autocorrelation in behavioral time series, we first modeled the residuals from the initial monotonic fit using an autoregressive moving-average (ARMA) process to determine the appropriate noise structure. This ARMA-informed residual model was then incorporated into the likelihood-based fitting procedure for all models. Model fit was assessed using Takeuchi’s Information Criterion (TIC), which accounts for both model complexity and potential misspecification, including remaining autocorrelation in the residuals. As behavioral time courses of target discrimination accuracy and subjective visibility showed similar theta-band rhythmic fluctuations regardless of valid and invalid trials across both left and right cueing conditions (Fig. 4), we plotted each participant’s TIC values of left cued valid / invalid trials and right cued valid / invalid trials for each of three model fit (19 participants x 4 trial combinations TIC values of each of the three models in Fig. 6). The damped oscillation model yielded a significantly lower TIC than both the monotonic decay model (mean L\TIC = 9.81 ± 0.63 for accuracy, 11.37 ± 0.64 for visibility) and the sustained oscillation model (mean L\TIC = 2.64 ± 0.29 for accuracy, 2.89 ± 0.35 for visibility) (Wilcoxon signed-rank test within subjects; Fig. 6), indicating a superior fit of the damped oscillation model to the empirical data. These results suggest that the third model, the damped oscillation model, best captures the temporal dynamics of behavioral accuracy and subjective visibility in retroperception.

**Fig. 6:**
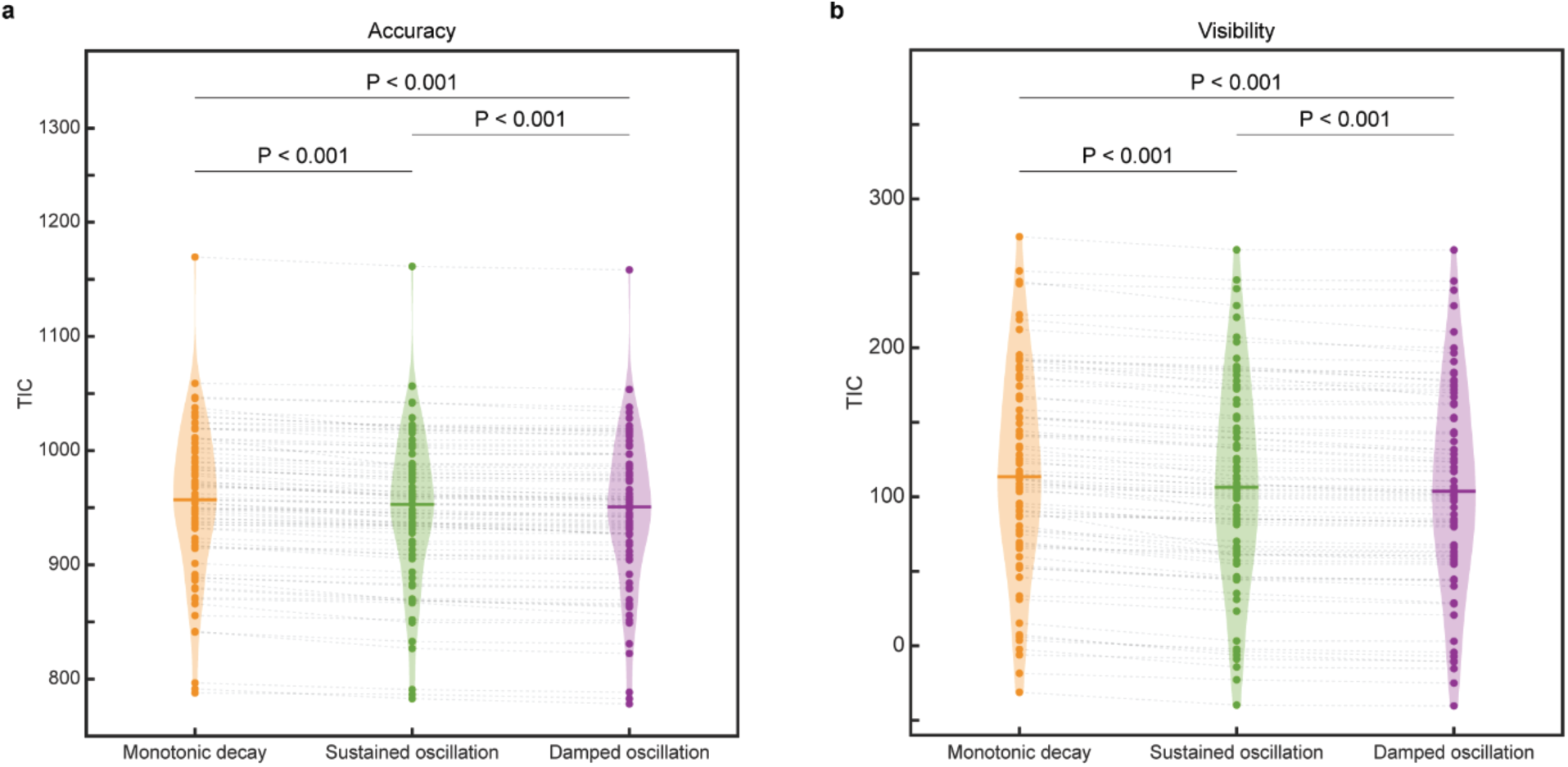
Comparison of Goodness-of-Fit across three models. Each colored dot represents the Takeuchi’s Information Criterion (TIC) value for one of the three models fitted to an individual temporal profile (orange, Monotonic decay model; green, Sustained oscillation model; purple, Damped oscillation model). As there are four possible trial combinations of retro-cue and its validity (left valid cue, left invalid cue, right valid cue, and right invalid cue) per participant, there are four TIC values for one of the three model fitted to four temporal profiles for behavioral accuracy and subjective visibility, respectively. Each of total 76 (19 participants x 4 trial combinations) TIC values from the three models are connected by gray dotted lines to illustrate within-subject model comparisons. Violin plots summarize the distribution of TIC values for each model with central horizontal lines indicating median values. Pairwise Wilcoxon sign-rank tests were used to assess statistical differences between models, with Bonferroni correction applied for multiple comparisons. **a** Model fits for target discrimination accuracy. The mean adjusted R^2^ scores were 0.06 ± 0.02 for the monotonic decay model, 0.18 ± 0.02 for the sustained oscillation model, and 0.21 ± 0.02 for the damped oscillation model. **b** Model fits for target visibility ratings. The mean adjusted R^2^ scores were 0.08 ± 0.02 for the monotonic decay model, 0.23 ± 0.02 for the sustained oscillation model, and 0.27 ± 0.02 for the damped oscillation model.

### Dynamic relationship between accuracy and visibility

Given the similar rhythmic fluctuations in both target discrimination accuracy and subjective visibility, we examined if their time courses are aligned with each other as previous study showed the positive correlation between subjectivity visibility and discrimination accuracy^7^. First, we examined if the IPFs of the time course of accuracy and those of the time course of visibility were the same or different. For each condition of the four possible combinations of cue-target position validity, the IPFs of discrimination accuracy time courses were not significantly different from those of subjective visibility time courses (Valid-Left*: t(18) = -0.82, p = 0.42*; Invalid-Left: *t(18) = -0.19, p = 0.85*; Valid-Right: *t(18) = -0.37, p = 0.72*; Invalid-Right: *t(18) = 0.01, p = 0.99*).

As both time courses of discrimination accuracy and subjective visibility showed theta-band frequencies as IPFs in all four conditions, we further investigated whether their time courses were in-phase at the theta-band frequencies. First, we detrended and filtered the time course of each of discrimination accuracy and subjective visibility using narrow band-pass filter spanning from lowest IPF to highest IPF (2.7 - 6.3 Hz). We then applied the Hilbert transform to each theta band-filtered time course to extract instantaneous phase at every time point. At every time point, the instantaneous phase difference between accuracy and visibility was calculated. We then performed the Rayleigh test to determine if these phase differences across all time points were uniformly distributed or clustered around a specific direction. We found that most participants showed the significantly consistent phase differences over time in accuracy and visibility in all four conditions (blue and red arrows in Fig. 7), except for one participant in the valid left-cued condition and two participants in the valid right-cued condition. Next, we performed a V test to determine whether these average phase angle differences were clustered around 0° across all participants. We further confirmed that the average phase angle differences were non-uniformly distributed around 0°, indicating robust phase alignment between accuracy and visibility at group level (Fig. 7).

**Fig. 7:**
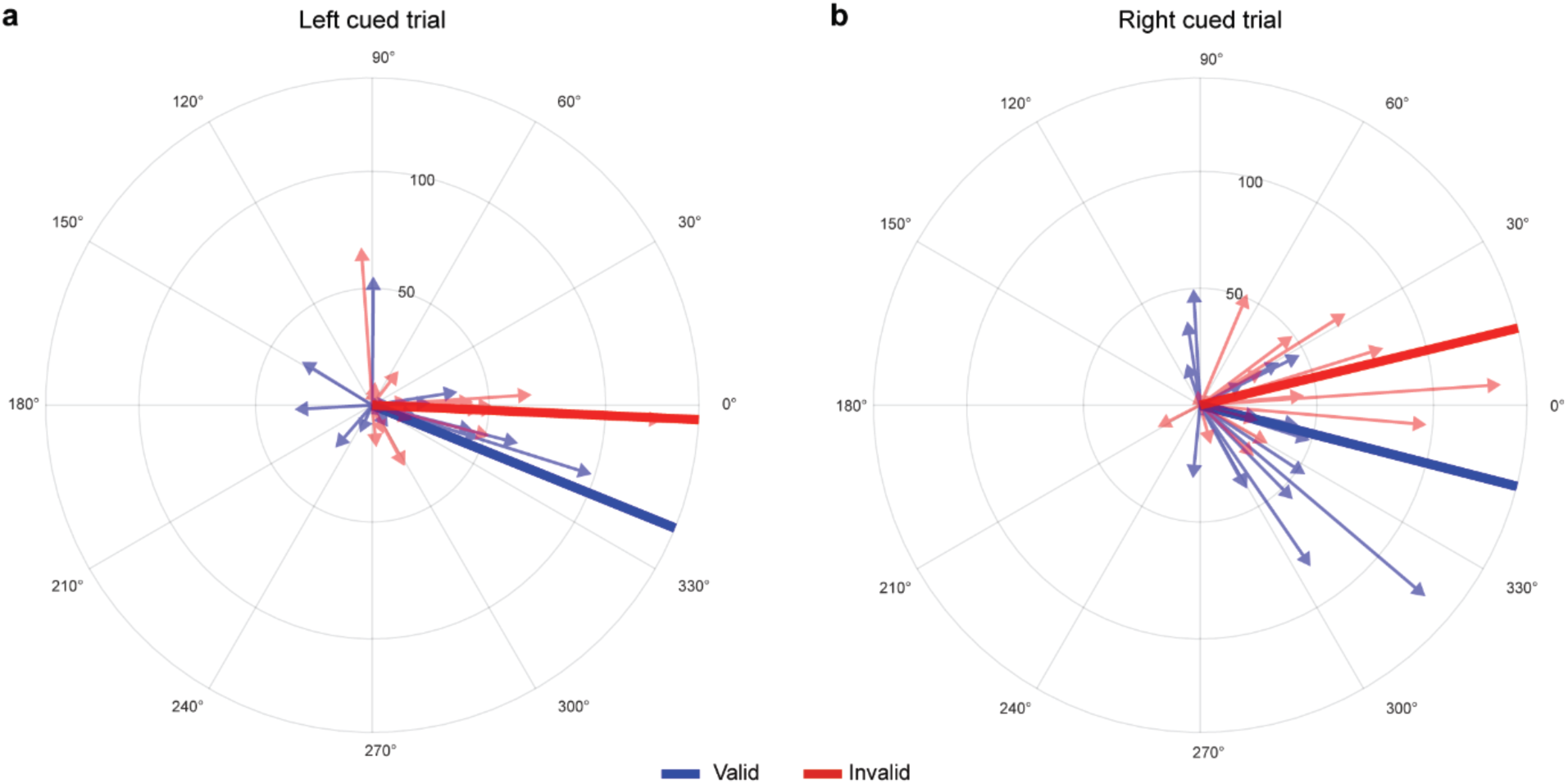
Phase alignment between discrimination accuracy and subjective visibility. Blue (red) arrows and lines represent valid (invalid) trials. Each arrow points to the average phase difference between theta-band filtered time course of accuracy and that of visibility for an individual participant, with arrow length indicating the strength of the phase clustering. The solid lines represent the average phase difference across all participants. a The average phase difference between the accuracy and visibility time courses, calculated across all time points in left-cued condition. In valid trials, 18 participants showed significantly consistent phase differences across all time points (blue arrows). One participant did not pass the Rayleigh test. At the group level, these average phase differences were non-uniformly distributed around 0° (V test; Z = 9.06, p = 0.001; mean difference = -22.05°; thick blue line). In invalid trials, 19 participants showed significantly consistent phase differences across all time points (red arrows). At the group level, these average phase differences were non-uniformly distributed around 0° (V test; Z = 13.06, p < 0.001; mean difference = -2.54°; thick red line). b The average phase difference between the accuracy and visibility time courses, calculated across all time points in right-cued condition. In valid trials, 17 participants showed significantly consistent phase differences across all time points (blue arrows). Two participants did not pass Rayleigh test. At the group level, these average phase differences were non-uniformly distributed around 0° (V test; Z = 9.60, p < 0.001; mean difference = -14.37°; thick blue line). In invalid trials, 19 participants showed significantly consistent phase differences across all time points (red arrows). At the group level, these average phase differences were non-uniformly distributed around 0° (V test; Z = 13.00, p < 0.001; mean difference = 13.61°; thick red line).

## Discussion

In this study, we investigated whether retroperception (i.e., the retrospective emergence of perceptual awareness) exhibits intrinsic rhythmicity and, if so, what temporal structure governs it. Using a dense-sampling retro-cue paradigm combined with both orientation discrimination task and its visibility rating, we showed that retroperception followed a pattern of damped oscillations at the theta-band frequency (-4 Hz) superimposed on an overall monotonic decay over time. These monotonically decaying damped oscillations were observed in both temporal profiles of the discrimination accuracy and the subjective visibility. Spectral decomposition using IRASA confirmed that these fluctuations reflect true rhythmic components rather than background noise fluctuations. Furthermore, model comparison analysis demonstrated that a damped oscillation model outperformed both monotonic decay model and sustained oscillation model in explaining the temporal dynamics of retroperception. Finally, the phase alignment between accuracy and visibility rating suggests that attention and perception are rhythmically aligned in the post-stimulus window. Taken together, these findings offer a unified framework in which conscious access operates not as a continuous stream, but as a temporally structured and rhythmically constrained process.

One previous study speculated on the rhythmicity in the retro-cue conditions but failed to observe it empirically^13^. This is likely due to the fact that the task-irrelevant but salient flash of four dots was used as a retro-cue, creating a classic backward masking effect^21^ on the preceding non-salient target and interfering with conscious perception. This backward masking caused a significant drop in target detection accuracy, making it impossible to reveal rhythmic structure in the retro-cue condition. However, current study used a brief dimming of a placeholder as a cue to avoid a typical masking effect on the target Gabor grating. This resulted in an average performance of 80% in the retro-cue condition, with no significant dips in the temporal profile of discrimination accuracy and subjective visibility, thereby allowing us to examine the rhythmic fluctuation in conscious access to past stimulus. As a result, we observed theta-band rhythmic fluctuation in retroperception under both left and right cueing conditions. This finding aligns with previous studies reporting that when attentional resources are distributed across both left and right hemifields, sampling frequency became identical across hemifields^19,22^. In our study, the dimming cue was relatively low in salience and exhibited 50% cue validity, thereby preventing attentional resources from being anchored to a single hemifield for an extended duration. Therefore, presenting a cue to either left or right visual field yielded similar peak frequencies, reflecting the distributed attentional resources across hemifields.

Previous studies of rhythmic attentional sampling have identified theta-band fluctuations in target sampling performance by resetting attention using salient cues prior to target presentation^13,17,18^. Even if no reset event occurred, spatially selective attention alternated between two locations prior to target presentation^22^. These previous findings suggest that the ongoing attentional sampling before target presentation alternates between the two locations at 4 Hz. In the present study, we extended the temporal window to 1.5 sec after the stimulus was gone, and found that attentional sampling was maintained and still alternated periodically at 4 Hz. Taken together, current study indicates that attentional sampling alternates between the two locations at theta-band frequencies before target onset and persists after target offset, further supporting the rhythmic attentional sampling hypothesis^17^.

The fact that the time course of subjective visibility also showed rhythmic fluctuation in the theta-band and was aligned in phase with the time course of target discrimination accuracy in the theta-band indicates a functional interaction between attention and perception in retro-cue condition. Specifically, unnoticed sensory information may be reactivated and consciously accessed by directing retrospective attention to the stimulus’s prior location^7–9^. Accordingly, target visibility was enhanced when the retro-cue was presented at a favorable phase of attention’s intrinsic rhythm. In contrast, cueing during an unfavorable phase diminished this enhancement. This rhythmic interplay likely reflects interactions between occipital sensory regions and fronto-parietal network^23–25^. Several theories propose that the occipital cortex samples sensory information rhythmically, while the attentional network modulates this sampling and superimposes its intrinsic rhythms^17,18,26,27^. Our findings support this framework, highlighting rhythmic synchrony between attentional processing and perceptual processing.

The observed temporal structure of retroperception is characterized not only by theta-band frequency oscillations but also by a gradual damping of their amplitude over time. While latent sensory trace can be reactivated by retro-cueing at a favorable phase of attentional sampling, the magnitude of reactivated information decreases over time. Therefore, there exists a temporal limit beyond which retro-cueing, even at an optimal phase, fails to reamplify the decaying sensory trace to above the critical threshold to gate it into conscious access. Beyond the rhythmic fluctuations, we observed overall decaying trend in both accuracy and visibility, consistent with the degradation of sensory traces in the absence of reactivation^28–31^. The overall decaying trend suggests that, even when retro-cueing occurs at an optimal phase and successfully reactivates the residual sensory trace, conscious access may still fail if the baseline signal has already fallen below a critical threshold. These trends are often overlooked in previous studies on pre-cued attentional sampling, which typically detrended behavioral data to emphasize only oscillatory components^14,18,32^. In contrast, our approach preserved the realistic time-dependent structure of retroperception by integrating damped rhythmic sampling with an overall decaying trend, highlighting the importance of considering baseline dynamics in future investigations of retroperception. These two distinct characteristics of retroperception reveal the limitations in the temporal flexibility of conscious access. For the unconscious sensory representation to be reactivated into consciousness, attention should be employed at its favorable phase and within the time window which sensory traces have not been decayed below a critical threshold. Therefore, our damped oscillation model, unifying attentional sampling with retroperception, provides rhythmically structured and temporally constrained window for conscious reactivation and imposes computational constraints on theories of consciousness.

Despite the strengths of our design, several limitations should be noted. Our experimental design incorporated both pre-cue and retro-cue conditions as well as cue validity manipulations; together with dense sampling across an extended CTI, this reduced the number of trials per condition. To boost trial counts per bin, we employed a 100 ms sliding time window, which functioned as a low-pass filter. Consequently, our spectral analyses were restricted to frequencies below 10 Hz, limiting our ability to examine higher frequency dynamics.

Several studies have proposed that rhythmic fluctuations in attention and perception at the behavioral level reflect underlying neural rhythms^16,33,34^. However, most of these studies have focused on pre-stimulus behavioral fluctuations in pre-cueing paradigm^14,18,35–37^, leaving post-stimulus rhythmic fluctuations relatively underexplored in retro-cue paradigm. Given that brain rhythms persist beyond stimulus offset^38^, behavioral fluctuations driven by neural oscillations are expected to extend into the post-stimulus period^39–42^. While our finding that retrospective attention gates conscious access to past visual stimulus in a rhythmically decaying fashion is largely consistent with those previous studies, future work on the underlying neural mechanism could strengthen the view that neural oscillations drive behavioral dynamics. Such studies could employ electroencephalography or magnetoencephalography during retro-cue paradigms to identify the electrophysiological signatures of retroperception and to directly test for damped oscillatory dynamics in neural activity.

In conclusion, our findings reveal that conscious access is neither an instantaneous nor uniformly sustained process, but one that unfolds within a temporally dynamic and intrinsically rhythmic window. Retroperception, long viewed as evidence of the mind’s capacity to reach back in time, is here shown to be governed by damped theta-band oscillations-periodic pulses of accessibility that gradually diminish as sensory traces decay. This rhythmic reactivation of latent information not only demonstrates that attention and perception continue to interact well beyond stimulus offset, but also delineates the temporal boundaries within which conscious access can become desynchronized from the initial sensory event. By capturing both the oscillatory and decaying components of this process, our study advances a temporally nuanced framework of conscious awareness-one that recognizes its remarkable flexibility, yet acknowledges its inherent temporal constraints.

## Materials and Methods

### Participants

A behavioral experiment was conducted with human participants in a dark room, as detailed below. Twenty-eight observers were initially recruited. Nine observers were excluded for excessive rejected trials caused by eye movements, leaving nineteen participants for analysis (mean age 22.9 ± 2.32, ten females, nine males). Detailed exclusion criteria are outlined in the Analysis section. All observers had normal or corrected-to-normal visual acuity and were naive to the purpose of the experiment. The observers gave informed consent to participate as paid volunteers. The study protocol was approved by the committee of the Institutional Review Board of the Korea National Institute for Bioethics Policy (P01-201909-13-004).

### Stimuli

All visual stimuli and experimental procedures were presented using Psychophysics Toolbox^43,44^ within MATLAB (Mathworks Inc.) running on a Mac mini. The observers viewed the 22” CRT (View Sonic PF817) monitor setting to a 100 Hz refresh rate and a resolution of 1,024 x 768 pixels. The monitor luminance was adjusted to generate a linear and constant mean luminance of 22 cd/m2. The viewing distance between the observers and the monitor was 60 cm. The observer’s head was stabilized using a chin rest. An eye-tracking camera (EyeLink 2000, SR research) was used to monitor observers’ eye movements throughout the experiment. The experimental paradigm of our study was adapted from Sergent et al.^7^. A uniform grey background was always on screen. Three stimuli simultaneously appeared in the beginning of each trial. A small bull’s-eye fixation (0.6° of visual angle, black) appeared at the center of screen and two circle placeholders (2.4° of visual angle, black) were aligned horizontally at 4 ° eccentricity to the left or the right of fixation. The fixation point remained visible until the visibility-response display and the placeholders stayed until the end of each trial.

A Gabor grating (2° of visual angle) was presented as the target for 50 ms within one of the placeholders. The spatial frequency of the grating was 2 cycles per degree of visual angle with a full width at half maximum of 1°. The grating’s orientation was randomly selected for each trial, either between 30 and 60 degrees or between 120 and 150 degrees. The grating’s average luminance matched the background luminance. The Michelson contrast of the grating was adjusted individually for each observer using a staircase procedure, aiming for approximately 80% accuracy on the orientation-discrimination task, excluding the effects of cueing. A flicker cue was the color change of the left or right placeholder from black to dark gray for 50ms.

Two response-screens were given for each trial. In the orientation-response screen, two small Gabor gratings (1° of visual angle, 4 cycles per degree, 0.5° full width half maximum) were presented at 1° eccentricities above and below fixation, both at maximal contrast. One of the gratings had the same orientation as the target and the other grating’s orientation was orthogonal to the target. Observers were required to indicate which of the two response gratings corresponded to the target’s orientation by pressing one of two response buttons within 1.5 seconds (’z’ key for the upper grating, ‘/’ key for the lower grating).

In the visibility-response screen, observers rated their subjective visibility of the target (1 to 8). The black vertical bar (1°x4°) was displayed at the center of screen, with eight equidistant positions along the bar representing visibility levels, ranging from ‘Not seen (1)’ to ‘Maximal visibility (8)’. The visibility-rating scheme was based on previous studies ^45,46^. Specifically, participants were instructed to respond 1 when they did not see a Gabor patch at all, respond 2-4 when they had a vague impression of a Gabor patch depending on the precision of this impression, respond 5-7 when they have seen a Gabor patch depending on the clarity and precision of their target perception, and respond 8 when they saw the clearest possible Gabor patch. A small white cursor (1°x0.5°) was initially located randomly at one of the positions on the vertical bar. Observers moved the cursor to their chosen visibility level using buttons mapped to the up or down directions and confirmed their selection by pressing the space key within 10 seconds.

### Experimental Procedure

Prior to the main experiment, an 80-trial staircase procedure^47,48^ was conducted to determine the target’s contrast level, using the same paradigm as the main experiment but without the cueing manipulation. This procedure was repeated twice for practice. The contrast was adjusted to a level at which the observer could barely discriminate the target’s orientation with 80% accuracy. The lowest contrast level from one of the staircase tasks was selected as the target contrast for the entire experiment.

In the main experiment, each trial began with a fixation screen that lasted for a random duration between 500 ms and 900 ms. A flicker cue was presented at a random time either between 1,550 ms and 100 ms before target onset (pre-cue condition; Fig. 2b) or between 100 ms and 1,550 ms after target onset (retro-cue condition; Fig. 2c). The inter-stimulus interval (ISI) was fixed within a range of 50 ms to 1,500 ms for both cue conditions, and data analysis was conducted based on the ISI.

The low-contrast target appeared randomly either on the same side as the cue (valid condition) or on the opposite side (invalid condition). The delay periods between the last stimulus onset (target for pre-cue condition and cue for retro-cue condition) and the onset of the response screen were randomly chosen between 150 ms and 450 ms.

Two response screens were presented with the time limits of 1,500ms and 10,000ms, respectively. The main experiment consisted of 984 trials, divided into six blocks of 164 trials each. In each block, target-present trials were equally divided between pre-cue and retro-cue conditions. Approximately 10% of the trials were catch trials, in which no target was presented. Eye calibration tasks were performed twice in a block, once at the beginning of the block and another in the middle of the block.

### Behavioral analysis

Analysis was conducted using MATLAB, Fieldtrip toolbox^49^, and CircStat toolbox^50^. Accuracy of the orientation discrimination task and subjective visibility ratings were analyzed as a function of the CTI for each cue timing (pre-cue vs. retro-cue) and cue validity (valid vs. invalid). At each time point t, we calculated accuracy by dividing the number of correct trials by the total number of trials within a 100-ms time bin, extending from t-50 ms to t+50 ms. To obtain accuracy for the subsequent time point we shifted the time bin by 10 ms and repeated this process throughout the entire CTI. In parallel, we calculated the subjective visibility of the target within the same 100 ms time bins.

We rejected trials if a participant’s gaze point deviated by more than 1° horizontally, because the target was presented exclusively on the horizontal plane relative to the fixation point. Following this procedure, 7 ± 1% of all trials were excluded from the analysis. Participants with missing values in any time bin due to rejected trials were excluded from further analysis.

### Spectral analysis using lRASA

Given that we applied a 100 ms sliding time window, which acts as a low-pass filter, we restricted our spectral analysis to frequencies below 10 Hz. To isolate rhythmic oscillation components and remove arrhythmic fractal fluctuations in the temporal dynamics of accuracy and visibility, we applied the Irregular-Resampling Auto-Spectral Analysis (IRASA)^20^ method in the Fieldtrip toolbox. We detrended each temporal dynamic using a quadratic function, applied a Hanning window, and zero-padded the data to 10 s for high frequency resolution. The 1/f fractal background activity was approximated using a moving time window of 75% of total length of temporal dynamic with a 50 ms step size. We defined an oscillatory peak as the strongest distinct peak exceeding the average 1/f background activity within 2.7-10 Hz, ensuring reliable estimates from at least four cycles in the temporal dynamic.

### Model fitting and comparison

We developed three models to determine which best explained the observed temporal dynamics of visual attention and awareness. Monotonic decay model: A second-order polynomial without oscillatory components (Equation (1)). Sustained oscillation model: A quadratic model with a first-order Fourier series added (Equation (2)). Damped oscillation model: A quadratic model with a linearly damping first-order Fourier series (Equation (3)). First, the monotonic decay model was fitted to the temporal dynamics using the lsqcurvefit function in MATLAB. The residuals from this initial fit were modeled using an autoregressive moving-average (ARMA) model to determine appropriate ARMA order. Subsequently, the determined ARMA model was employed in maximum likelihood estimation fitting for all three models. For the sustained oscillation model and the damped oscillation models, the monotonic decay model fitted to the data was treated as the fixed trend. We calculated the adjusted R^2^ score and Takeuchi’s Information Criterion (TIC) to compare the relative quality of the models, considering both goodness-of-fit and model complexity. Pairwise Wilcoxon test was used to assess the statistical differences of goodness-of-fit between each model, with Bonferroni correction applied for multiple comparisons. A commonly accepted guideline suggests that a difference of approximately 2 units in the TIC value indicates a meaningful improvement in model fit quality^51^.

### Frequency and Phase alignment

To assess alignment between the temporal dynamic of accuracy and the temporal dynamic of visibility, we examined their IPFs and instantaneous phases. We subtracted the visibility IPF from the accuracy IPF under the corresponding cue conditions; we then tested these IPF differences using a t-test.

To assess phase alignment, we used the Hilbert transform on detrended and band-pass filtered temporal dynamics (2.7-6.3 Hz). We then subtracted the instantaneous phase of visibility from the instantaneous phase of target discrimination accuracy at each corresponding time point. We tested the resulting phase differences of each individual for clustering using the Rayleigh test in the CircStat toolbox. After applying false discovery rate (FDR) correction using the Benjamini-Hochberg (BH) method^52^, we identified significant clusters of phase difference for each participant. Finally, we tested whether the individual phase difference was significantly clustered around 0° using a V test, which is a Rayleigh test with a specific target direction.

