## Supplementary Figure 1 for "Retrospective attention reveals a decaying theta rhythm in conscious access to a preceding stimulus"

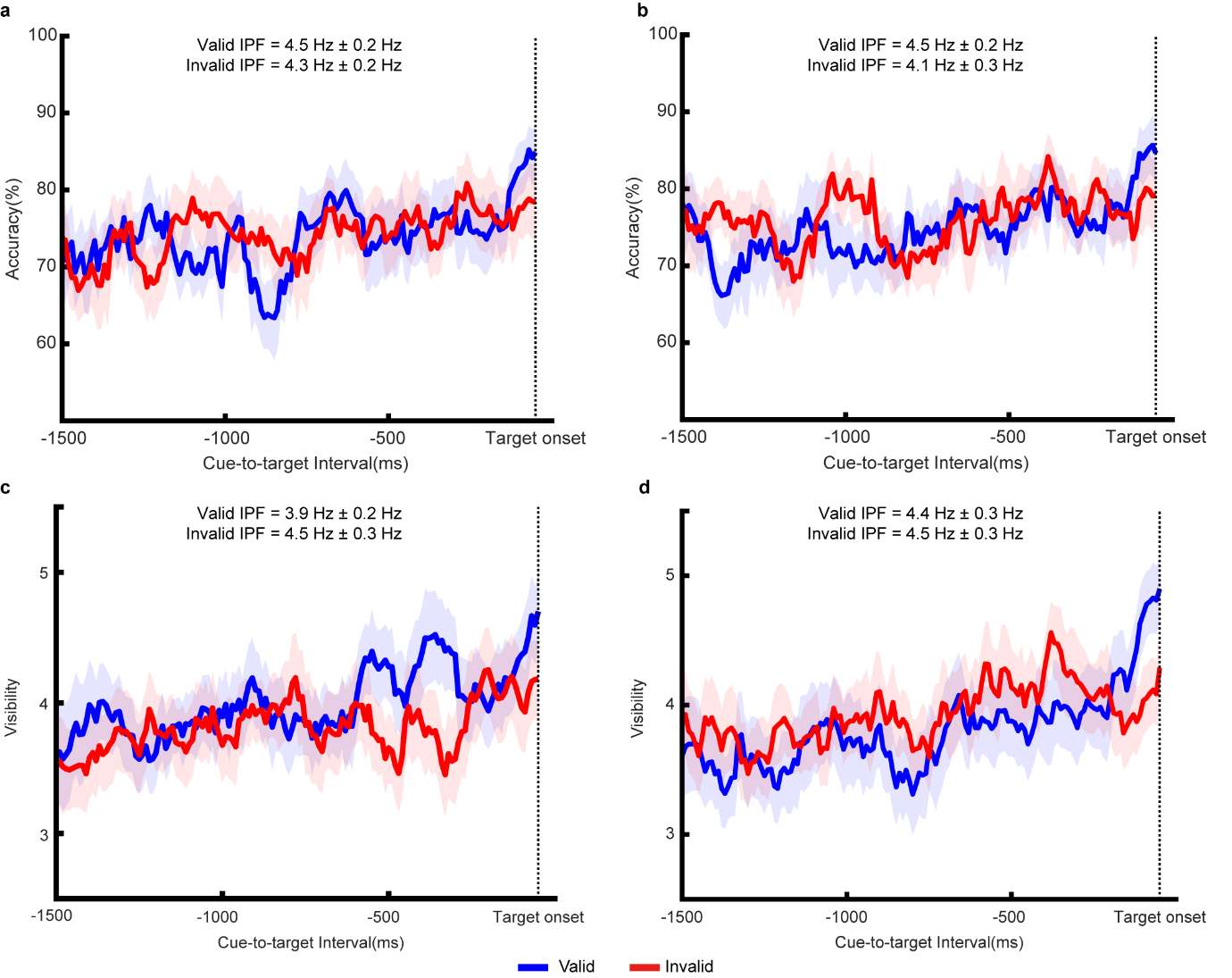

**Fig. S1: Temporal dynamics of visual performance and awareness in pre-cue conditions.**

Black dotted lines mark the time of the target onset. Blue (red) lines and shaded areas correspond to valid (invalid) cue trials. Solid lines indicate the average performance or visibility rating across participants (n = 19), shaded areas represent the SEM.

**a**, Target discrimination performance as a function of CTI for left-cued trials. The mean IPF in valid trials was 4.5 $\pm$ 0.2 Hz, compared with 4.3 $\pm$ 0.2 Hz in invalid trials.

**b**, Target discrimination performance as a function of CTI for right-cued trials. The mean IPF in valid trials was 4.5 $\pm$ 0.2 Hz, compared with 4.1 $\pm$ 0.3 Hz in invalid trials.

**c**, Perceived visibility of the target as a function of CTI for left-cued trials. The mean IPF in valid trials was 3.9 $\pm$ 0.2 Hz, compared with 4.5 $\pm$ 0.3 Hz in invalid trials.

**d**, Perceived visibility of the target as a function of CTI for right-cued trials. The mean IPF in valid trials was 4.4 $\pm$ 0.3 Hz, compared with 4.5 $\pm$ 0.3 Hz in invalid trials.
